# Plasma microRNAs as predictive biomarkers of liver fibrosis and cirrhosis in patients with HBV-associated disease

**DOI:** 10.1101/2020.07.21.213546

**Authors:** Tie-Zheng Wang, Dong-Dong Lin, Yong-Hong Zhang, Yue-Wei Zhang, Ning Li

## Abstract

Both pathogenesis and detection of liver fibrosis and early cirrhosis are elusive. Our study focused on role of microRNAs (miRNAs) as predictive biomarkers of liver fibrosis and early cirrhosis. Methods: We recruited patients with chronic hepatitis B background and divided them into three subgroups according to the pathological staging results: no liver fibrosis group (CHB 11, 12.0%), liver fibrosis group but without liver cirrhosis (LF 41, 44.6%), and early liver cirrhosis group (LC 40, 43.4%). Results: Levels of miR-122 in the CHB and LF group were significantly elevated compared with the LC group. Meanwhile, upregulation levels of miR-146a-5p, miR-29c-3p, miR-223 were seen in the CHB group versus LC. MiR-122-5p was best able to differentiate between patients with LC and without LC (area under the curve (AUC) 0.734). It can also distinguish patients with LF from those without LF (AUC 0.681). Panel including miR-122-5p miR-29c-3p and miR-381-3p resulted in an AUC of 0.700 in detection of LF. Meanwhile panel including miR-122-5p, miR-29c-3p, MiR-223 turned out to be good predictors, resulted in an AUC of 0.752 to discriminate LC in HBV-associated disease. HBV load, ALT and AST showed significant positive correlation with miR-122-5p and miR-381-3p. Conclusions: Our study explored changes of plasm miRNA levels in HBV-associated disease patients, constructed diagnostic miRNAs panel to predict liver fibrosis and early cirrhosis. Meanwhile In the process of liver fibrosis and cirrhosis, there are involving not only common pathway molecules, but also its specific pathway molecules.

## Introduction

The most common cause of liver fibrosis is chronic viral hepatitis B in china. 30% of cirrhosis cases are caused by to hepatitis B worldwild [1] . Although liver biopsy remains the golden criterion standard for staging of fibrosis, this procedure is invasive, painful and carries the risk of complications[2]. In order to overcome the limitations of liver biopsy, a number of non-invasive techniques have been investigated for the assessment of liver fibrosis and cirrhosis[3]. Current noninvasive measures are nonspecific and insensitive. Thus, a sensitive, specific, and noninvasive tool, reflecting the early pathogenesis of liver fibrosis or cirrhosis before the advent of severe complications, is required.

MicroRNAs (miRNAs) may play this role in the future. miRNAs are a series of short noncoding regulating RNA molecules that contain 20–25 nucleotides, bind to the 3’-untranslated region of target miRNAs, and interferes with gene expression at the post-translational level by way of inducing translational arrest, which in turn reduces or prevents protein synthesis[4]. Substantial evidence implicates miRNA involvement not only in maintaining normal liver function but also in liver disease processes. For example, circulating levels of liver-specific miR-122 have been detected upregulated in viral, alcoholic, metabolic, and drug-induced liver injury [5–7]. Moreover, the evidence of the correlation between the expression of miRNAs and liver fibrosis has increased. Consequently, these brought miRNAs to the frontline in the development of noninvasive diagnostic tools.

In the present study, we selected 7 candidate miRNAs, miR-122-5p, miR-21-5p, miR-146a-5p, miR-29c-3p, miR-223, miR-22-3p, MiR-381-3p, based on our previous study [8]. We tried to use reverse transcription-quantitative polymerase chain reaction (RT-qPCR) to quantify plasm miRNAs in patients with HBV infectious background. We hypothesized that these 7 candidate liver-specific miRNAs would be detected and showed significant differences among patients with CHB (F0), liver fibrosis (F1-F3) and early cirrhosis (F4). MiRNA panel were used to improve the diagnostic ability of liver fibrosis and cirrhosis.

## Materials and methods

### Clinical specimens. All patients were selected based on clinical reports

All patients were examined at You’an Hospital Hospital, from October 2016 to September 2017. Enrolled patients were for HBsAg positive, and did not have any other liver disease such as hepatitis C, cholangiocarcinoma, or autoimmune, alcoholic and metabolic liver diseases. Written informed consent was obtained from each of the participants and the study was approved by the Ethics Review Committee (ERC) of Beijing You’an Hospital. The study was conducted in accordance with the principles of the Declaration of Helsinki, the standards of Good Clinical Practice (as defined by the International Conference on Harmonization).

### Liver histology

Specimens in the study were collected through two methods: ultrasound-guided percutaneous liver biopsy or wedge biopsy during surgery. Metavir fibrosis-scoring [9] was applied to determine the severity of the fibrosis. F0 refers to no fibrosis and F4 refers to early liver cirrhosis. This procedure was performed by two senior pathologists independently.

### Noninvasive assessment of liver fibrosis

The aspartate aminotransferase-to-platelet ratio index (APRI) and fibrosis-4 (FIB-4) scores : they were two formulas derived from routine blood markers and calculated as follows: APRI model, APRI=AST (ULN) × 100/PLT[10]; FIB-4 model, FIB-4=(AGE × AST) / PLT × (ALT1/2)[11].

### Transient elasticity (TE)

FibroScan® 502 (Elsevier, France) was used for the transient elasticity imager. Patient was supine and the right arm was lifted to fully expose the intercostal space. The right anterior axillary line was selected to the 7th, 8th and 9th intercostal spaces of the axillary line for continuous detection. The mean value was the final Liver stiffness measurement (LSM) result.

### RNA Extraction and quantification

Vein blood samples were collected using the BD Vacutainer Plus and centrifuged at 3,000 rpm for 15 minutes within four hours after collection. The supernatant plasma was immediately recovered and stored at −80°C. Plasma miRNAs were extracted using a total RNA isolation kit (QuantoBio). Synthetic Caenorhabditis elegans miRNA (cel-miR-67) was used as an exogenous control and added to the plasma lysate before extraction. The quantity of miRNAs was determined by SYBR-based qRT-PCR, according to manufacturer’s instructions (Quantobio). Escherichia coli polyA polymerase was used to add a polyA tail at the 3′ end of the RNA molecules. With oligo (dT) annealing, a universal tag was attached to the 3′ end of the cDNAs during cDNA synthesis using reverse transcriptase (Quantobio). Quantitative PCR was performed with miRNA-specific forward primers and a universal reverse primer mix according to the universal tag. The relative expression level of miRNA was calculated using the 2^−ΔΔct^.

### Experimental design and statistical analysis

All data were analyzed using SPSS 20.0 statistical software (SPSS, Chicago, IL, USA) and GraphPad Prism 5 software (GraphPad software, La Jolla, CA, USA). Measurement data were expressed as mean ± standard deviation. The differences between normally distributed numeric variables were evaluated using Student’s t-test, HBV RNA levels were lg transformed to enable parametric statistical tests. Non-normally distributed variables such as data of these seven miRNAs were analyzed by Mann–Whitney U-test. Spearman correlation analysis was used to visualize the relationship between the HBV viral load & liver function parameters and the seven miRNAs. The performance of plasma miRNAs was examined by the area under the corresponding receiver–operating characteristic (ROC) curve (AUC). Logistic regression were carried out to construct diagnostic miRNAs panel. All *P*-values were two-sided, and with P < 0.05 considered significantly different.

## Results

### Patient Characteristics

A total of 92 patients were enrolled in this study. The mean age was 40.58 ± 11.43 years (range: 16-60 years). BMI was 22.87 ± 3.04 kg/m^2^ and the MELD index was 5.25 ± 4.40. Among these patients, eleven patients were in F0 stage, 16 patients were in F1 stage, 12 patients were in F2 stage, thirteen patients were in F3 stage, and 40 patients were in F4 stage. Patients were divided into three subgroups by liver histology results: CHB (F0), LF (F1-3) and LC (F4). The characteristics of participants were presented in Table 1. Age, MELD, LSM, FIB-4, Serum ALT, GGT, ALP and HBV DNA levels were significantly different among subgroups (P<0.05). There were no significant difference in the distribution of Gender, Child-Pugh, APRI, BMI, AST among these three groups (P>0.05).

**TABLE 1.**
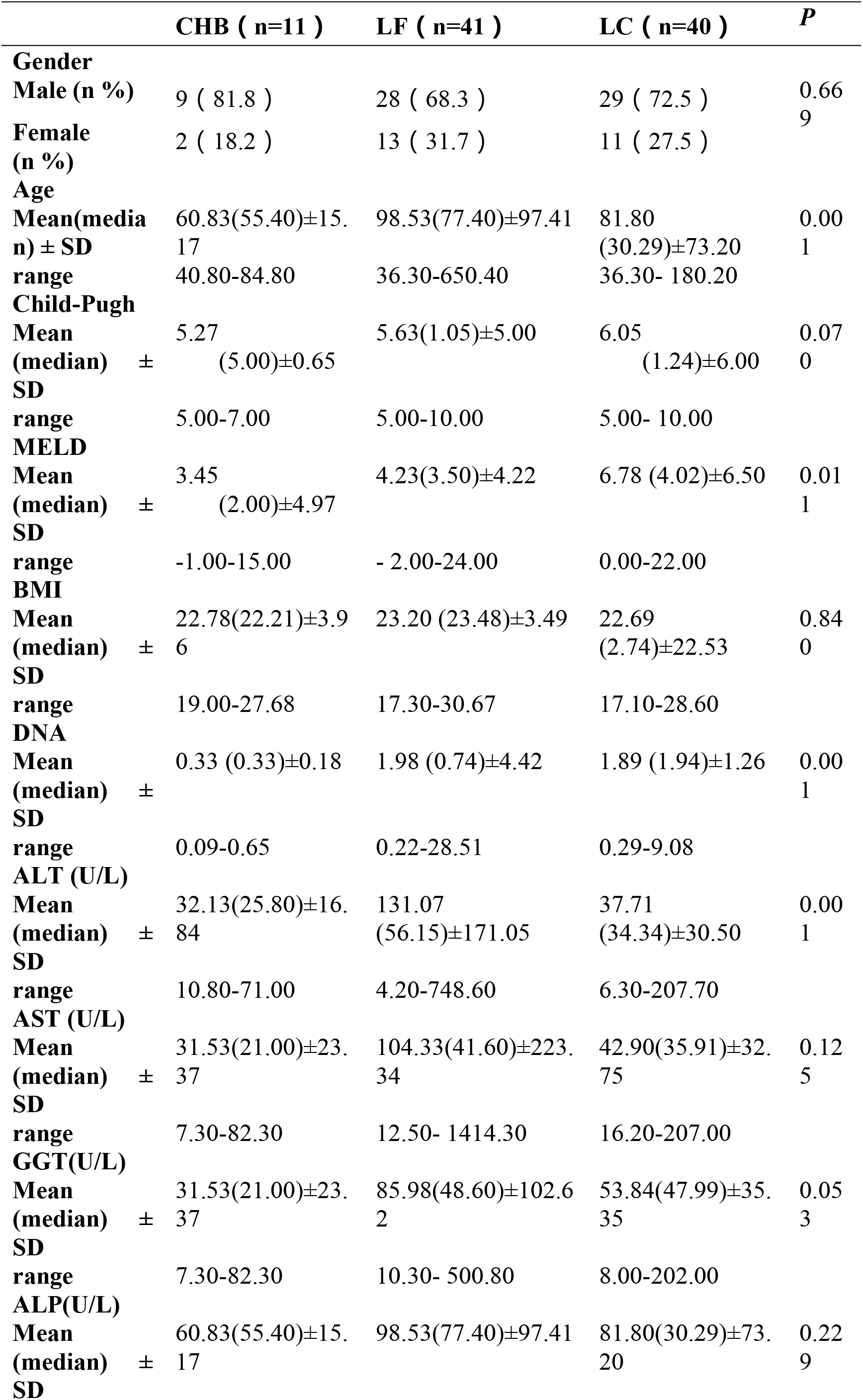
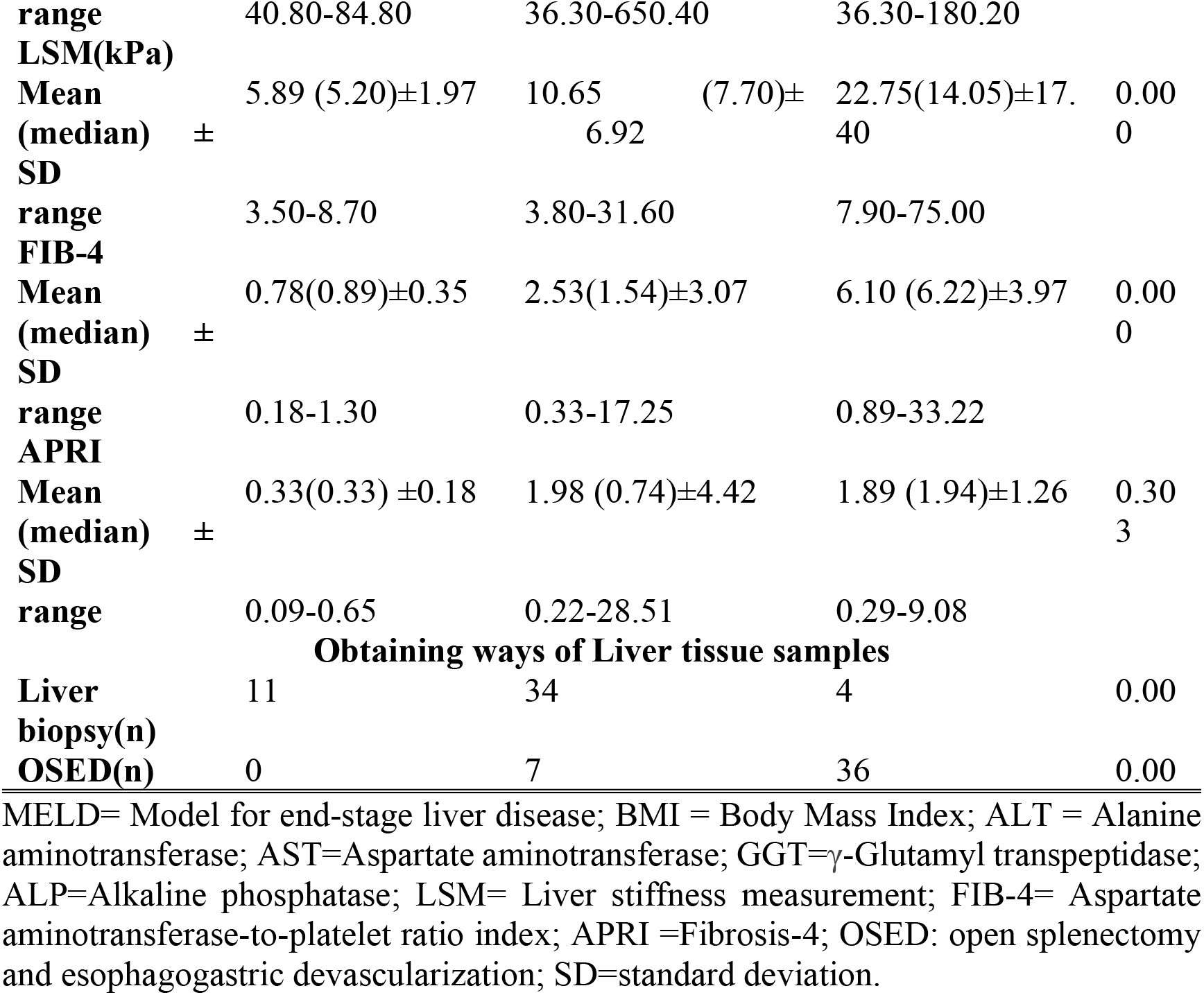
Demographic and clinical characteristics of the CHB LF and LC groups

#### Relative expression Levels of 7 Candidate miRNAs

Figure 1 shows the relative plasm levels of the 7 candidate miRNAs determined by RT-qPCR and normalized to Synthetic Caenorhabditis elegans miRNA (cel-miR-67). Plasma levels of miR-122 were significantly elevated both in CHB and LF group compared with LC group (P = 0.028; P = 0.000 respectively). Mean while, an increase in the level of miR-146a-5p miR-29c-3p, miR-223 was seen in the CHB group versus LC (P = 0.037; P = 0.007; P = 0.017 respectively). None significant difference was observed for miR-21-5p, miR-22-3p, and miR-381-3p in our study.

**FIGURE 1.**
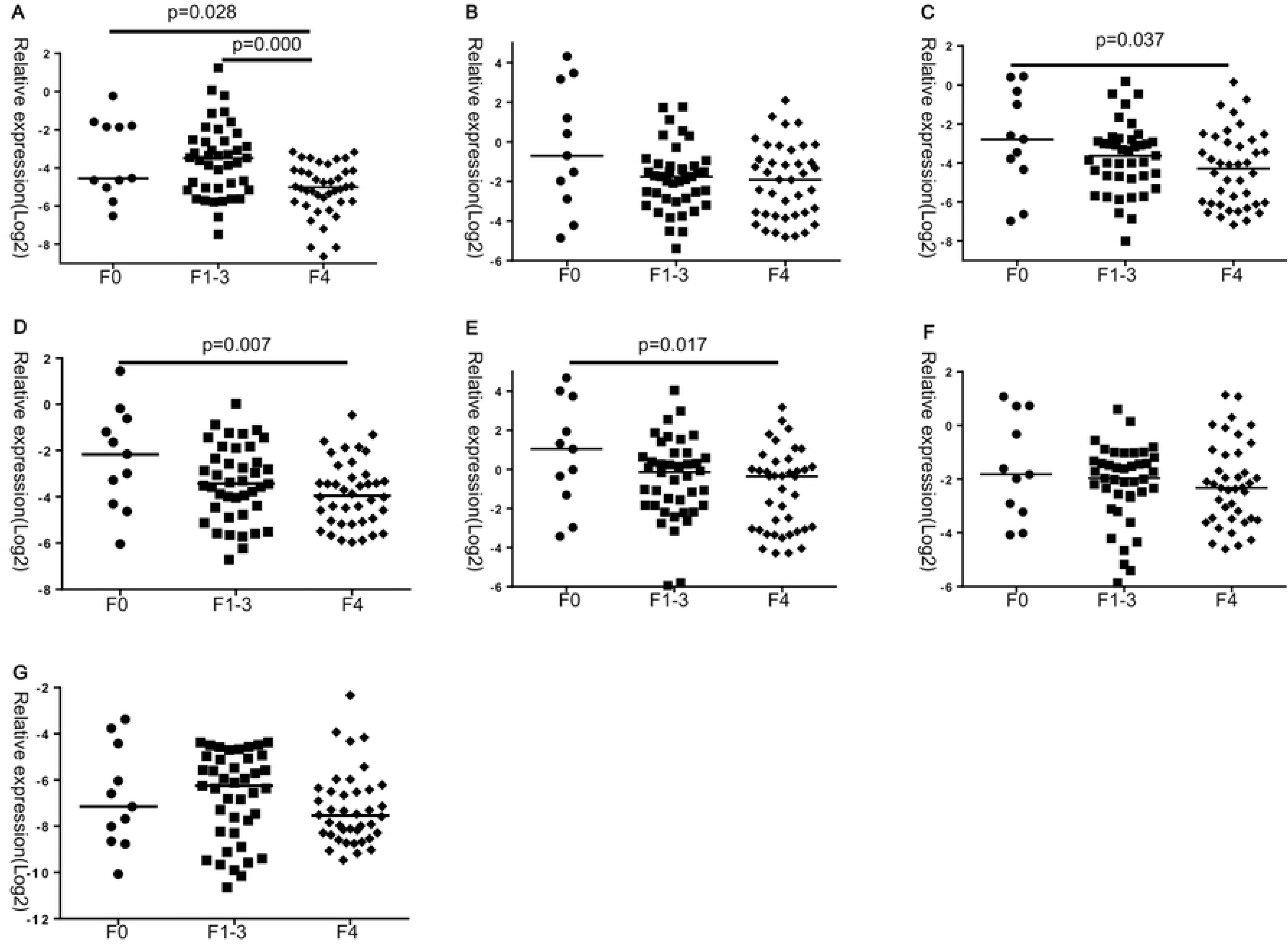
Plasm miRNA levels in CHB (F0) group, LF (F1-3) group, and LC (F4) Group. (A) miR-122-5p, (B) miR-21-5p, (C) miR-146a-5p, (D) miR-29c-3p, (E) miR223, (F) miR-22-3p, and (G) miR-381-3p. Lines on scatter plots represent median.

### Distinguishing Liver fibrosis and early LC from HBV-associated disease

ROC curve analysis was used to determine the ability of individual miRNA to distinguish patients with LF (F1-3) from patients with CHB (F0) and LC (F4). Or LC (F4) from LF (F1-3) and CHB (F0). MiR-122-5p was best able to differentiate between patients with LC and those with LF and CHB (area under the curve [AUC] 0.734; 95% confidence interval [CI] 0.634-0.834; P =0.000) (Fig.2A), and distinguish patients with LF from those with CHB and LC. (AUC 0.681; 95% CI0.570–0.793; P=0.003) (Fig.2B).

**FIGURE 2.**
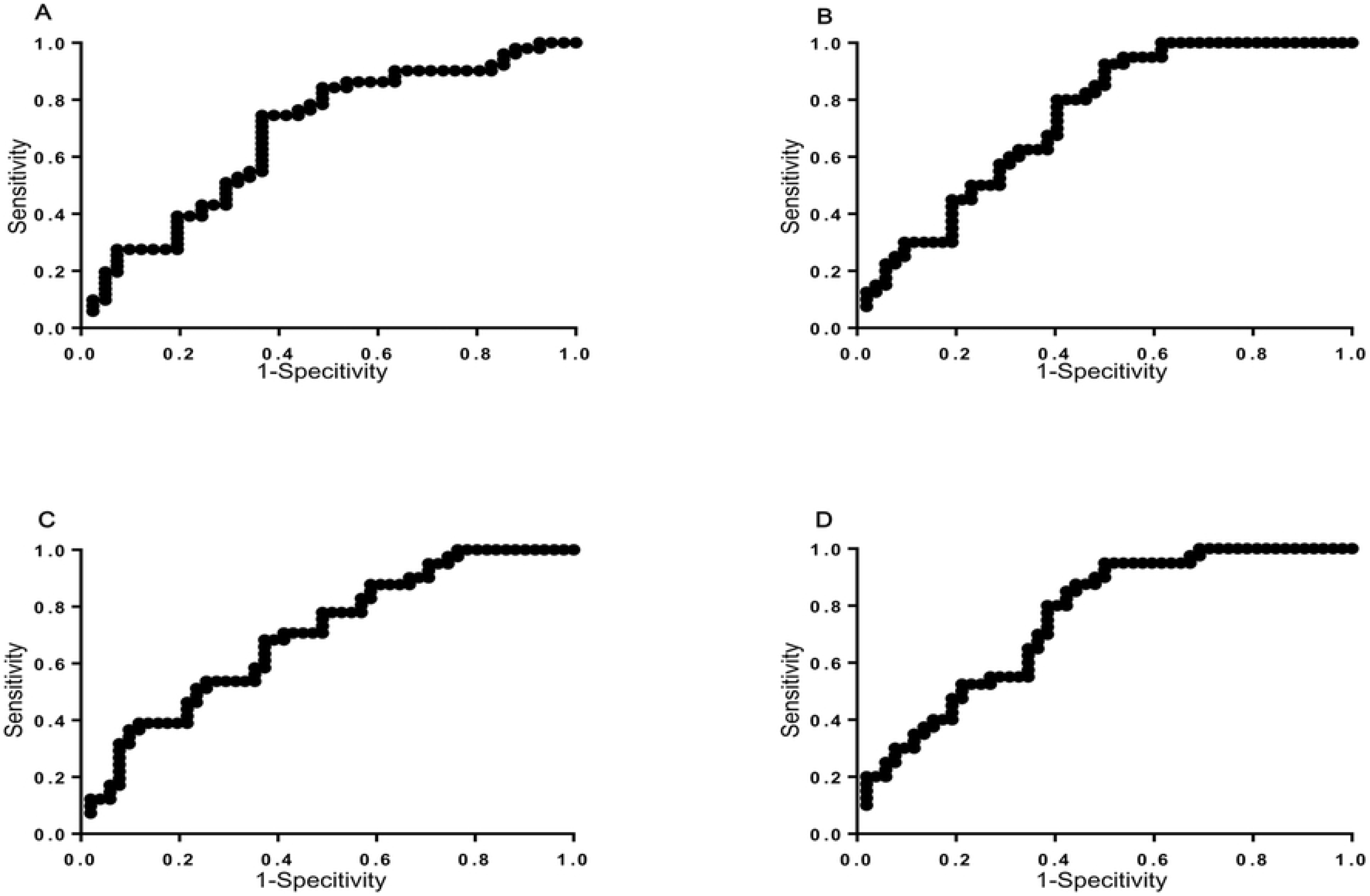
(A).ROC curve for plasma miR-122-5p level to discriminate patients with LF (F1-3). The area under the curve is 0.681(95% CI 0.570–0.793; P=0.003). (B).ROC curve for plasma miR-122-5p level to discriminate patients with LC (F4). The area under the curve is 0.734 (95% CI 0.634–0.834; P≤0.001). (C).ROC curve for the combination of miR-122-5p miR-29c-3p and miR-381-3p plasma levels to discriminate patients with LF (F1-3). The area under the curve is 0.700 (95% CI 0.596–0.791; P≤0.001). (D).ROC curve for the combination of miR-122-5p, miR-29c-3p and miR-223 plasma levels to discriminate patients with LC (F4). The area under the curve is 0.752 (95% CI 0.651–0.836; P≤0.001). CI=confidence interval; miR=microRNA; ROC=receiver operating characteristic.

### Diagnostic performance of predictive and noninvasive panel combined with selected plasma miRNAs

Logistic regression analysis revealed that panel 1, including miR-122-5p miR-29c-3p and miR-381-3p, resulted in an AUC of 0.700 (sensitivity 68.29 %, specificity 62.75 %, PPV 59.6%, NPV 71.1 % CI 0.596–0.791; P =0.0002) in the detection of LF in HBV-associated disease patients (Fig.2C).

Logit (P) =1.277-0.530*miR-122-5p+0.300* miR-29c-3p −0.046*miR-381-3p. Panel 2,including miR-122-5p,miR-29c-3p,miR-223 turned out to be good predictors ,resulted in an AUC of 0.752 (sensitivity95.00%, specificity50.00%, PPV59.4%, NPV92.9 % . 95% CI 0.651–0.836; P<0.0001) to discriminate LC in HBV-associated disease(Fig.2D).

Logit (P) =−2.105+0.660*miR-122-5p-0.342* miR-29c-3p+0.248*miR-223.

### Correlation of plasm miRNA Levels with HBV viral load and Liver Function

Spearman rank correlations were performed to determine whether plasm miRNA levels were associated with viral load and liver function in patients with HBV-associated disease. HBV load, ALT and AST showed significant positive correlation with miR-122-5p (*r*=0.500, *P*=0.000; *r*=0.458, *P*=0.000; *r*=0.242, *P*=0.021 respectively) and miR-381-3p (*r*=0.554, *P*=0.000; *r*=0.242, *P*=0.021; *r*=0.239, *P*=0.022 respectively).Neither GGT nor ALP showed any significant correlation with any of the miRNAs. (Table 2). We also explored the correlation of plasm miRNA Levels with 3 kinds of current wildly used noninvasive diagnostic tools.LSM was found to correlate negatively with miR-122-5p, miR-146a-5p,miR-29c-3p,miR-223,miR-22-3p and miR-381-3p(r=−0.350, P=0.003; r=−0.290, P=0.014; r=−0.342,P=0.004; r=−−0.274, P=0.021; r=−0.260, P=0.028; r=−0.304,P=0.010 respectively),APRI showed negative correlation with miR-21-5p,miR-146a-5p,miR-29c-3p,miR-223 and miR-22-3p(r=−0.277, P=0.008; r=−0.363, P=−0.000; r=−0.225,P=0.031; r=−−0.414, P=0.000; r=−0.212, P=0.043 respectively). Mean while FIB-4 showed significant negative correlation with miR-122-5p, miR-21-5p,miR-146a-5p,miR-29c-3p,miR-223 and miR-22-3p (r=−0.419, P=0.000; r=−0.272, P=0.009; r=−0.376,P=0.000 respectively).

**TABLE 2.**
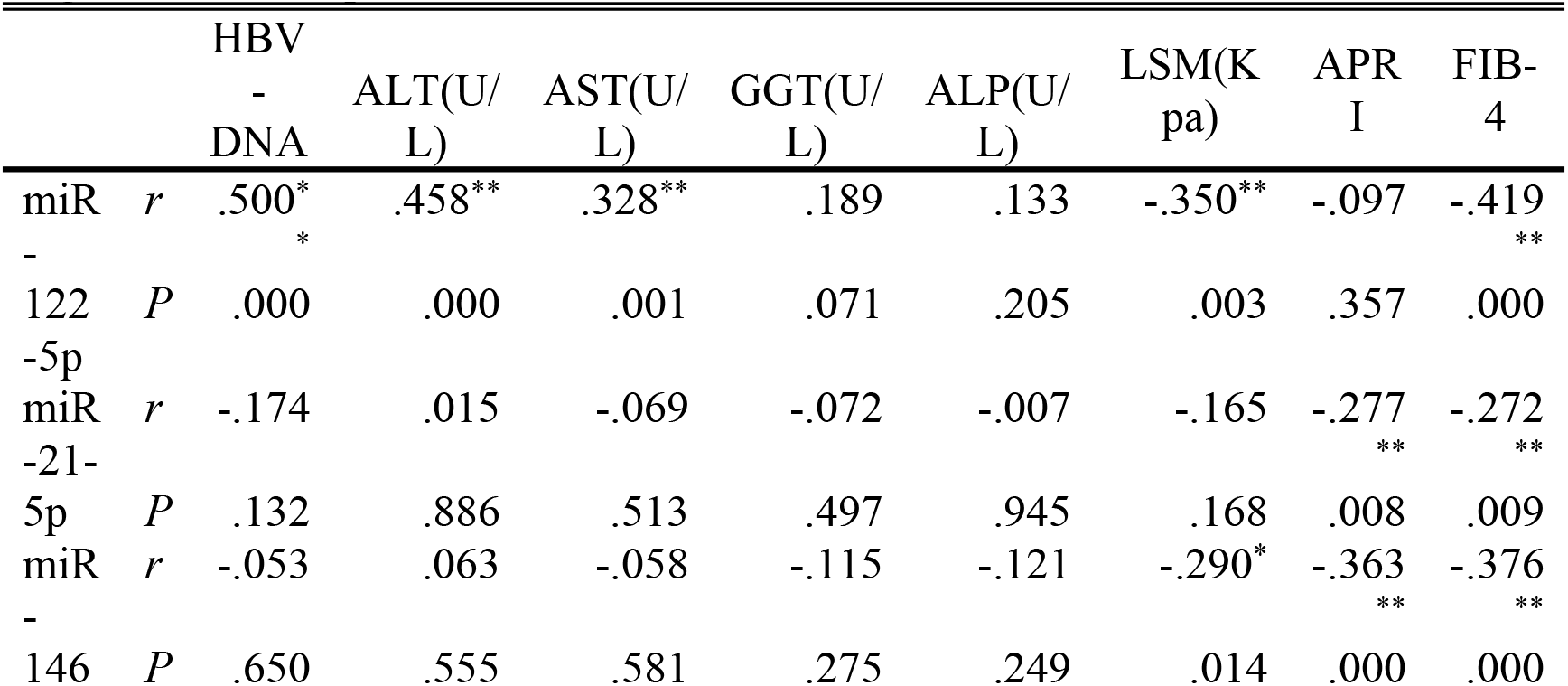
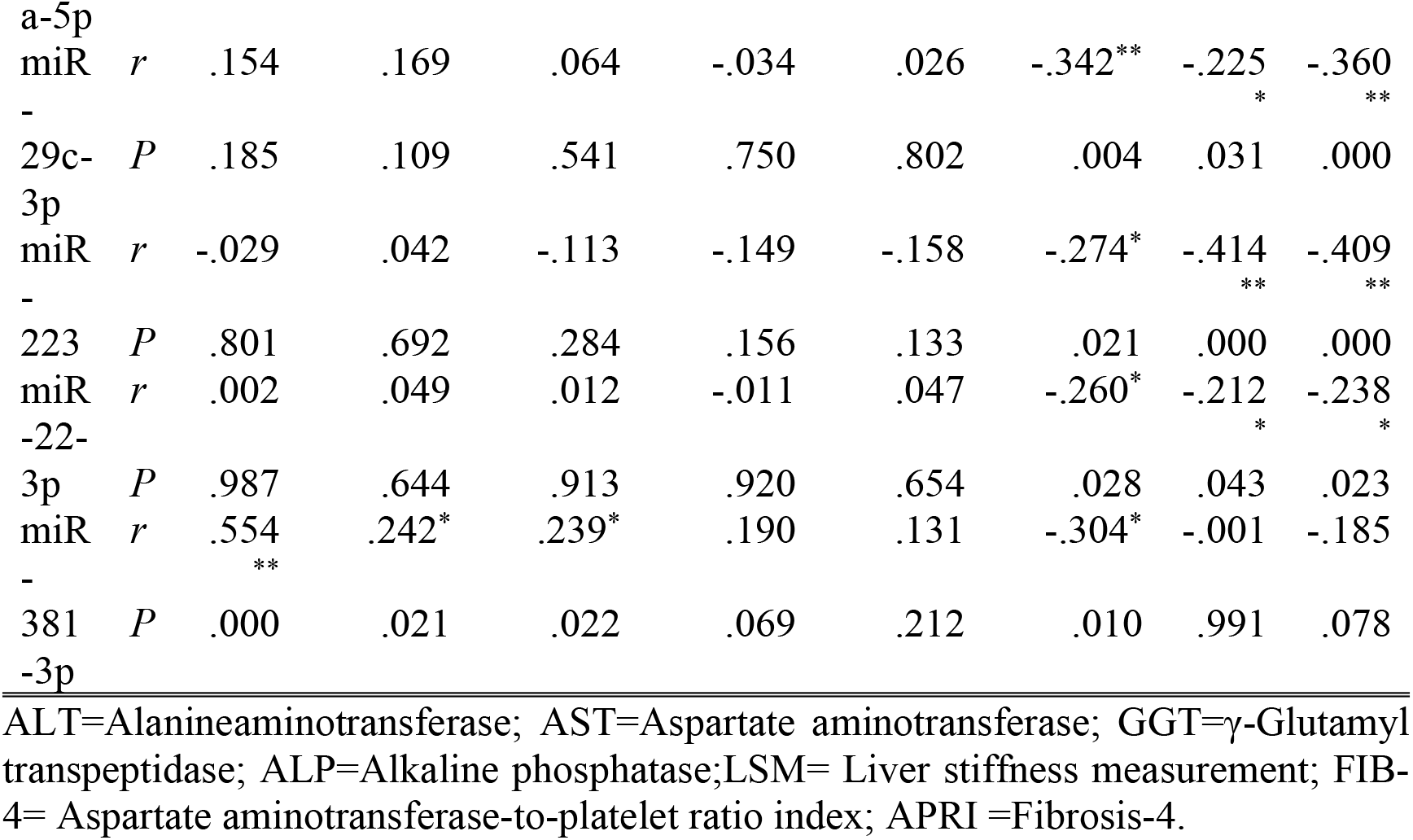
Correlation of miRNA levels with HBV load ,liver function and noninvasive diagnostic tools in patients with HBV-associated disease

## Discussion

Hepatitis B is the most common cause of liver fibrosis / Cirrhosis in china. Cirrhosis is the most advanced stage of fibrosis, which may develop into hepatocellular carcinoma (HCC). It can be said that liver cirrhosis is the final pathological stage of fibrosis, and fibrosis is the precursor of cirrhosis[12]. There is a need for noninvasive biomarkers to identify individuals at risk of developing into liver fibrosis and cirrhosis, which may guide the appropriate timing of antiviral therapy or pay much attention on surveillance of complications. Several reports suggested that different aetiologies of chronic liver disease may manifest a differential diagnosis levels of specific circulating miRNAs [13]. So we intentionally enrolled HBV-associated disease patients in this study to remove the interference factors. To our knowledge, CHB and Cirrhosis had been diagnosed by clinical experiences. During our routine clinical practice, we had found several clinically diagnosed CHB and cirrhosis patients were not consistent with their pathological results. We speculated that the liver itself has a strong compensatory capacity, so that some patients with early liver cirrhosis maybe clinical manifestations of mild or asymptomatic. Meanwhile some cases had been clinically diagnosed of cirrhosis, but the pathological changes were slight. So we believe that accurate clinicopathological staging is the basis for this study to obtain credible data. Specimens, in this study, were collected through two methods: ultrasound-guided percutaneous liver biopsy (n=49) and wedge biopsy (n=43) during surgery. The former mainly gather in F0-F3, whereas the latter gather in F4. METAVIR scoring systems, applied to determine the severity of the fibrosis, is the most commonly used histopathological systems to evaluate chronic hepatitis. In this scoring systems, F0 refers to no fibrosis and stage F4 refers to early liver cirrhosis. Totally 43 cases had been clinically diagnosed with liver cirrhosis and received surgical treatment. Pathological results in 7 cases (16.2%) (1 case of F1, 2 cases of F2, 4 cases of F3) had not reach the early diagnosis of liver cirrhosis (F4). The reason may be existing the chance that wedge biopsy during surgery were not of the accurate situation.

MiRNAs are abundant and very stable molecules in serum, plasma and liver tissue, regulating a diverse aspects of liver functions. There are several miRNAs having aberrant expressions and play a key role in the pathogenesis of liver fibrosis[14]. It was shown that altered miRNA patterns after chronic liver disease highly affect the progression of fibrosis by their potential to target the expression of extracellular matrix proteins and the synthesis of mediators of profibrogenic pathways. In addition, the deregulation of miRNAs may present long before the onset of the disease[15].

Circulating miRNAs, deriving from blood or body fluid, make them to be easily accessible potential biomarkers for the evaluation of liver fibrosis and Cirrhosis. Stability of concentrations, in whole blood at room temperature for up to 12 hours, is another advantage[16]. Although different status of circulating miRNAs, such as liver injury, inflammation and fibrosis, has there internal relation, the functional link between the regulation of miRNA levels and their roles in the specific cell compartments involved in fibrogenesis have remained unclear[17].

The capability of plasm miRNAs to detect early fibrogenesis and cirrhosis in HBV-associated disease patients was also assessed in the present study. The result of single profile is not particularly satisfactory. ROC analyses showed that only plasm miR-122 were effective predictive biomarker of LF (AUC=0.681; P=0.003) and LC (AUC= 0.734; P=0.000) in patients with HBV-associated disease.

MiR-122 miR-146a-5p miR-29c-3p miR-223 were used as candidate molecules for construction of the panel. Our strategy of incorporating miRNAs with the most significant differences between groups into the regression diagnostic panel yielded better diagnostic efficacy. By using a combination of 3 miRNAs (miR-122-5p miR-29c-3p and miR-381-3p/ miR-122-5p, miR-29c-3p, miR-223), we have demonstrated potential diagnostic use for discriminating LF /LC in HBV-associated disease patients. Constructing a regression diagnostic panel, can significantly improve the diagnostic efficacy of miRNAs, which maybe a research strategy on miRNA study. Based on this fact, we can draw a conclusion: In the process of liver fibrosis and cirrhosis, there are existing not only common pathway molecules (miR-122-5p, miR-29c-3p), but also its specific pathway molecules (miR-381-3p, miR-223).

In the study, three non-invasive techniques TE, APRI and FIB-4 undergone deep research. Most of these miRNAs showed significant negative correlations with noninvasive tools above. Show that there is an intrinsic relationship between miRNA and noninvasive examination.

In conclusion, the high stability of miRNAs in circulation makes them perfect biomarkers, especially for the diagnosis of early stage, asymptomatic HBV-associated diseases. So we believed that: the miRNA, especially circulation miRNA has a better prospect worthy of our in-depth study.

## Acknowledgments

This work was supported by grants from National S&T Major Project of China (2017ZX10203205) and National Grand Program on Key Infectious Diseases (2015ZX10004801)

## Author Contributions

**Conceptualization:** D-DL, NL

**Data curation: D-DL,NL.**

**Formal analysis: T-ZW.**

**Funding acquisition: D-DL.**

**Investigation: D-DL, NL.**

**Methodology: T-ZW, Y-HZ and Y-WZ.**

**Project administration: D-DL, NL.**

**Resources: T-ZW, Y-HZ and Y-WZ.**

**Software: T-ZW.**

**Supervision: D-DL, NL.**

**Validation: D-DL, NL.**

**Visualization: D-DL, NL.**

**Writing – original draft: D-DL, NL .T-ZW.**

**Writing – review & editing: D-DL, NL**

